# Decoding the spatiotemporal dynamic neural representation of repetitive facial expression imitation

**DOI:** 10.1101/2024.02.26.582020

**Authors:** Qi Liu, Xinqi Zhou, Siyu Zhu, Can Liu, Yanmiao Yang, Chunmei Lan, Xinwei Song, Benjamin Becker, Keith M. Kendrick, Weihua Zhao

**Affiliations:** The Center of Psychosomatic Medicine, Sichuan Provincial Center for Mental Health, Sichuan Provincial People’s Hospital, University of Electronic Science and Technology of China, Chengdu, 611731, China; Institute of Brain and Psychological Sciences, Sichuan Normal University, Chengdu, 610066, China; School of Sport Training, Chengdu Sport University, Chengdu, 610041, China; The State Key Laboratory of Brain and Cognitive Sciences, The University of Hong Kong, Hong Kong, 999077, China; The Clinical Hospital of Chengdu Brain Science Institute, MOE Key Laboratory for NeuroInformation, School of Life Science and Technology, University of Electronic Science and Technology of China, Chengdu, 611731, China

**Keywords:** Facial emotion imitation, spatiotemporal dynamics, neural representation, mirror neuron system, information flow

## Abstract

Imitating facial emotion expressions can facilitate social interactions, although the behavioral and neural spatiotemporal dynamics is unclear. Here participants (N=100) imitated facial emotions repeatedly over one month (16 times in total) with neural activity measured on three occasions using functional near-infrared spectroscopy. Additionally, the transfer effect of repeated imitation on emotional face and scene perception was assessed by fMRI with multivariate pattern analysis. Valence specific imitation performance was facilitated by the alterations in the similarity of spatiotemporal patterns evoked in the mirror neuron system (MNS) with information flow moving progressively towards the inferior frontal gyrus as the as the number of times of imitation increase. Furthermore, MNS representation predictive patterns of processing emotional faces, but not scenes, were enhanced. Overall, these findings provide a neural changes of information flow within MNS and advance our understanding of the spatiotemporal dynamics from novice to proficient of facial emotion imitation.

## Introduction

Imitation, refers to the ability to simultaneously observe and replicate an action displayed by others^1–3^ and represents a fundamental social process^4,5^ which serves to facilitate affiliation with others and helps vicarious learning^6^. Humans develop the ability over time to detect and recognize observed complex facial expressions via repetitive imitation or imitative learning over time^7^. The brain cortical mirror neuron system (MNS) comprising the inferior frontal gyrus (IFG), inferior parietal lobule (IPL) and superior temporal sulcus (STS) has been shown to play a critical role in synthesizing the observation and imitation of movements^8–11^ leading to a reconceptualization of the motor system from an entity involved not only in the imitation of movement but also as a highly complex system contributing to the perception and mapping of observed actions in humans supporting a range of social cognition functions^12–14^. Indeed, MNS dysfunctions have been implicated in empathic or imitation problems in a number of psychiatric disorders, especially in schizophrenia^15^ and autism spectrum disorder (ASD)^16^. A large number of studies have investigated the function of the MNS using motor or action imitation paradigms^17,18^ but in the context of daily social interactions the ability to express and recognize facial expressions is more critical^19^ and requires repetitive imitation or imitative learning^7^. In contrast to motor imitation, imitation of facial expression represents a global sensorimotor simulation of others’ emotions rather than a mere muscle-specific resonance (e.g. grasp)^8,20^. Thus, a fundamental issue addressed by the current study concerns how repeated imitation of face expressions influences imitative performance and what spatial and temporal encoding changes occurring within the MNS are predictive of this.

Within the MNS, imitation relies on a defined information flow, with initial visual input flowing from the STS to IPL and representing motoric description of the action and thence to the IFG involved processing the goal of the action. Subsequently, the imitative commands are fed back to the STS to allow matching between the sensory predictions of imitative motor plans and the visual description of the observed action^10^. Several patient-based studies have suggested that the IFG plays a more direct role in imitation^21–23^. However, few studies have examined this pathway in the context of repetitive facial expression imitation. Additionally, one recent study using functional near-infrared spectroscopy (fNIRS) found different roles for these three key MNS regions in imitation of positive and negative face emotion expressions^24^. However, whether emotion imitation dependent changes exhibit a valence-specific trajectory of changes during a period of repetitive imitation has not been established.

Here, the current study aimed to investigate four objectives: (1) to test whether individuals improved their face expression imitation performance across valences (positive vs. negative) following repetitive imitation (i.e. 16 times in one month); (2) to use fNIRS to investigate MNS pattern similarity changes across valences at three different time points during the period of repetitive imitation; (3) to determine valence-dependent alterations in specific pathways within the MNS regions across the three different time-points; (4) to use fMRI to confirm the transfer effect of repeated imitation on neural processing of observed face emotions by measuring MNS representational changes before and after repetitive imitation.

Thus, we leveraged a repetitive facial expression imitation task (FEI), combined with an objective facial expression recognition software (FaceReader, Noldus 7.1) to measure behavioral imitation performance^24^ across repeated sessions (n = 16, see **Figure 1**) and additionally MNS activity was recorded using fNIRS on three occasions (2^nd^, 9^th^ and 16^th^ FEI). Given that an increase in pattern similarity between trials within a condition over time would reflect an increased neural population overlap and integration after a period of consolidation^25–27^, we calculated MNS neural pattern similarity (NPS) across negative and positive facial expression imitation separately. Directed phase transfer entropy (dPTE) assessing information processing dynamics or effective connectivity in the MNS was employed to determine how its three key cortical regions interact or transfer information during repetitive FEI^28–31^. Furthermore, multivariate pattern analysis (MVPA)^32,33^ and informational connectivity^34–36^ were performed to evaluate whether the MNS changes during repetitive FEI also resulted in altered patterns of responses during observation of an independent set of emotional faces using task functional magnetic resonance imaging (fMRI) before and after the period of repetitive FEI. MVPA is a powerful method for establishing patterns of activation which are predictive of what emotions are being viewed and informational connectivity can synchronize changes in the presence of multivariate patterns over time to provide a more sensitive measure of functional connections between different regions^35,37^. We predicted that mirror neuron changes during the simulation of observed facial expressions would in turn facilitate patterns of neural encoding underlying perception and empathy towards others expressing emotions^38^. Overall, the present study has allowed us to understand for the first time how spatiotemporal changes in MNS processing during repeated imitation of positive and negative facial expressions that may improve social cognition and empathy by taking advantage of methodological advances in neuroimaging analyses. It is hoped that this progress may help to establish the utility of efficient imitation-based interventions for the individuals with social behavior dysfunction.

**Figure 1.**
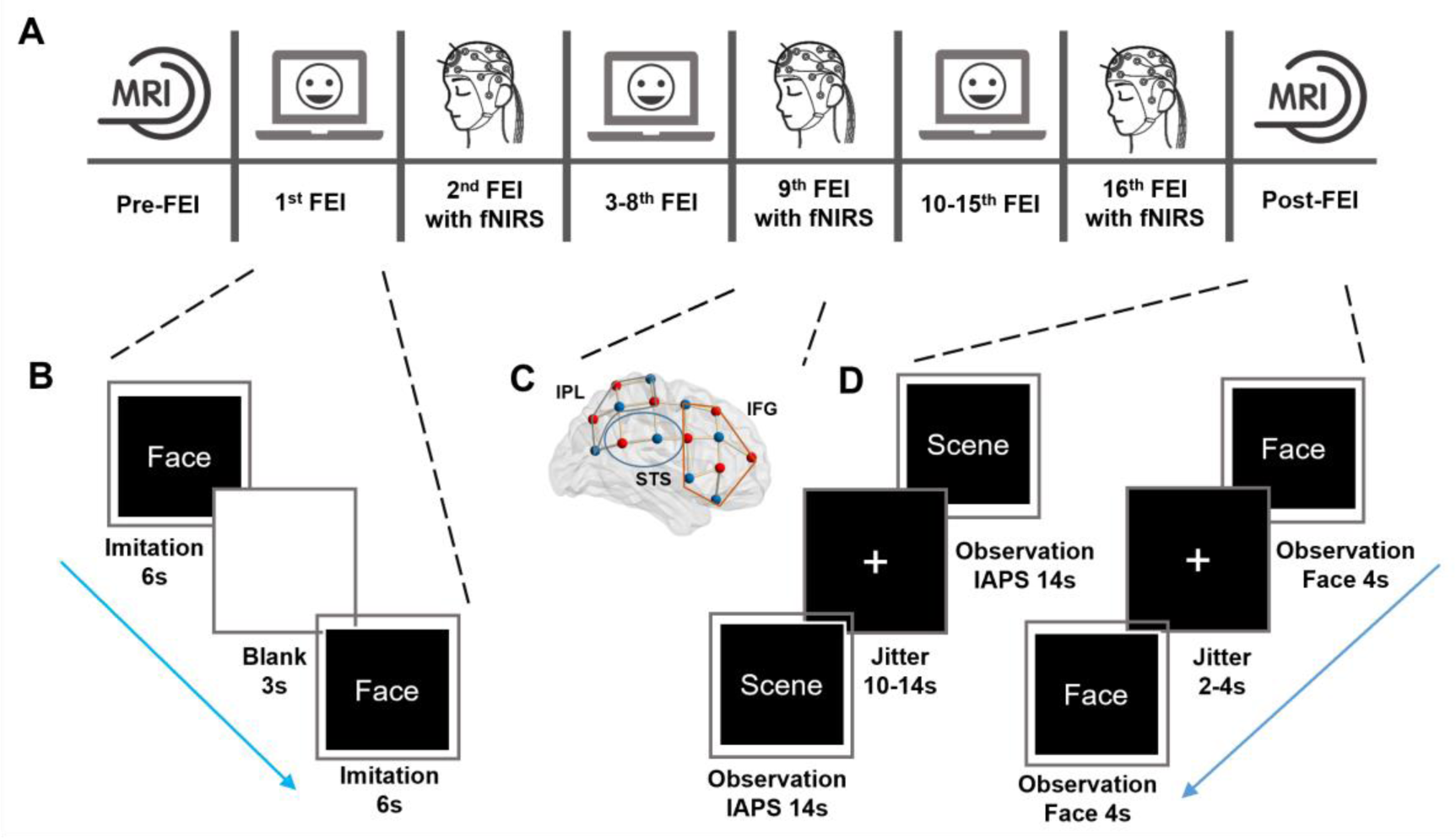
Experimental Protocol. (A) All participants (n = 80) were required to repetitive facial expression imitation (FEI) 16 times over one month (3-4 times per week) with three fixed fNIRS sessions (during 2^nd^, 9^th^ and 16^th^ FEI) and fMRI assessments during emotional face and scene perception paradigms before and after the repetitive FEIs. (B) FEI procedure. (C) The location of fNIRS opodes over the three key regions of the mirror neuron system (IFG = inferior frontal gyrus; STS = superior temporal sulcus; IPL = inferior parietal lobule). (D) Emotional face and scene perception (International Affective Picture System-IAPS) tasks during fMRI scans.

## Results

### Behavioral results

Considering the confounding effects of being unfamiliar with imitating expressions and the potential negative impact of wearing an fNIRS electrode cap on the accuracy of imitation performance, we only retained auto-decoded imitation performance data by Facereader (5.09Hz, see ***Methods***) from the 3^rd^-8^th^ (defined as the early phase FEIs) and 10^th^-15^th^ (defined as the late phase FEIs) repetitions. Two-way ANOVA revealed significant main effects of the number of repetitions (in total 12 times, *F*(11, 869) *=* 5.133, *p* < 0.001, *ƞ^2^_p_* = 0.686) and emotion valence (positive emotion vs. negative emotions, *F*(1, 79) *=* 172.79, *p* < 0.001, *ƞ^2^_p_* = 0.686), with greater area under the curve (AUC) for imitation of positive emotion (mean ± SD = 2.77 ± 1.25) compared to negative ones (mean ± SD = 0.98 ± 0.37). Exploratory statistical analyses also showed the AUC of imitating a positive face expression was significantly greater than that of imitating any of the negative ones (*F*(1, 79) *>* 83.39, *ps* < 0.001, **Table S1**). Notably, a significant interaction effect between emotion valence and the number of repetitions (*F*(11, 869) *=* 3.963, *p* < 0.001, *ƞ*^2^ = 0.061) was found. Repeated measures correlation analyses were conducted to explore the repetitive imitation effects (see ***Methods***). There were significant positive correlations between the AUC of positive imitation performance and the overall number of repetitions during the whole FEIs (*r*_rm_(879) = 0.179, *p*_FDR_ < 0.001, 95% confidence intervals (CI) = [0.115, 0.242]) due primarily to the early phase repetitions of FEIs (*r*_rm_(399) = 0.170, *p*_FDR_ = 0.003, 95% CI = [0.073, 0.236]) but not the late phase ones (*r*_rm_(399) = 0.055, *p* = 0.271, 95% CI = [−0.043, 0.152]). On the contrary, it was only negatively associated with the number of repetitions for negative FEIs during the late phase repetitions (*r*_rm_(399) = −0.158, *p*_FDR_ = 0.004, 95% CI = [−0.253, −0.062]). For each individual correlation (see ***Methods***, **Figure 2A** top), greater within-subject correlation was found between the AUC for imitation performance and the number of repetitions in positive relative to negative emotion expressions (positive: mean rho = 0.165 ± 0.053, negative: mean rho = −0.051 ± 0.047; *t*(79) = 3.121, *p* = 0.003, **Figure 2A** bottom).

**Figure 2.**
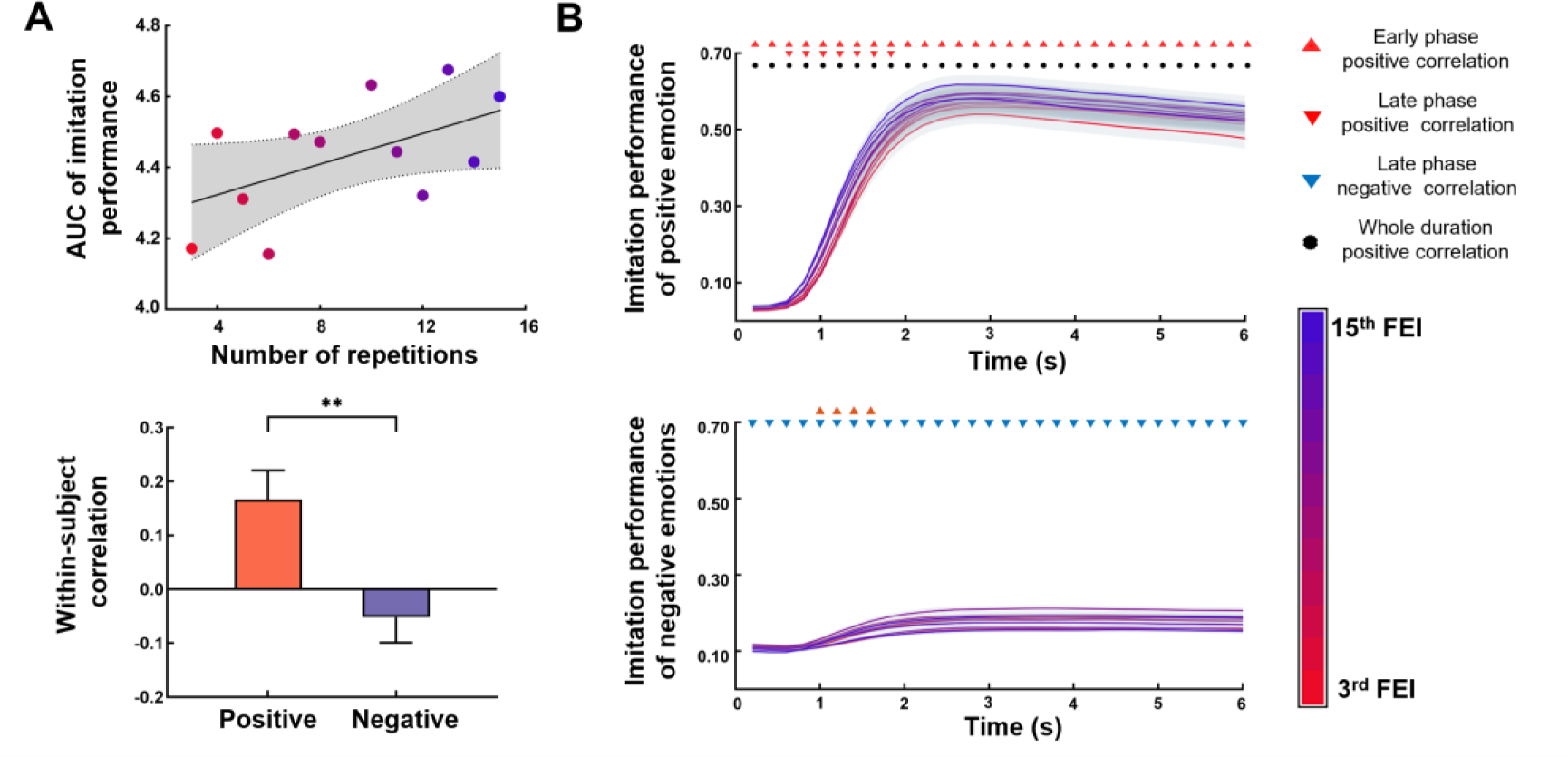
Behavioral results of repetitive facial expression imitation. (A) Top: The correlation between the area under the curve (AUC) of imitation performance and the number of repetitions for an example subject. Bottom: Within-subject correlation for imitation of positive and negative emotions. ** *p* < 0.01. (B) Time series analyses for imitation performance of imitating positive (top) and negative (bottom) emotions. Red upper triangles indicate that imitation performance and the number of repetitions were significantly positively correlated in early phase FEIs. Red lower triangles indicate positive correlations in late phase FEIs. Blue lower triangles represent negative correlations in late phase FEIs. Black dots represent positive correlations in the whole duration of FEIs. Error bands represent standard errors. All correlations were calculated by repeated measures correlation and *p* values were FDR-corrected. AUC = area under the curve; FEI = facial expression imitation; Early phase = 3^rd^ – 8^th^ FEI; Late phase = 10^th^ – 15^th^ FEI; Whole duration = early and late phases.

Further time series analyses indicated that positive imitation performance during the whole imitation process (0-6s) was positively correlated with the overall number of FEI repetitions (*p_FDR_ <* 0.001) and during the early phase (*p*_FDR_ < 0.031), but only at 0.6-1.8s (*p*_FDR_ < 0.035) during the late phase FEIs (**Figure 2B** top). For negative emotions, significant positive correlations occurred for early phase FEIs (1-1.8s, *p*_FDR_ < 0.031) and negative ones for late phase FEIs (0-6s, *p*_FDR_ < 0.016, **Figure 2B** bottom). Subsequently, two-way ANOVA for the onset-time of successful imitation (see ***Methods***) revealed significant main effects of the number of repetitions (in total 12 times, *F*(11, 869) *=* 1.827, *p* = 0.046, *ƞ^2^_p_* = 0.023) and emotion valence (positive emotion vs. negative emotions, *F*(1, 79) *=* 99.890, *p* < 0.001, *ƞ^2^_p_* = 0.558), with faster onset-time for successful imitation of positive emotion (mean ± SD = 2.19 ± 0.52s) compared to negative emotions (mean ± SD = 2.87 ± 1.30s). The repeated measures analysis showed a significant negative correlation between the onset-time of successful imitation of positive emotion and the number of repetitions during the overall period of FEIs (*r*_rm_(879) = −0.105, *p*_FDR_ = 0.003, 95% CI = [−0.170, −0.040]) and in the late phase FEIs (*r*_rm_(399) = −0.120, *p*_FDR_ = 0.032, 95% CI = [−0.215, −0.022]) but not in the early phase FEIs (*p* = 0.196). We did not find any correlations for imitating negative emotions (*p*s > 0.065). Overall, these results indicated that early phase FEIs selectively improved the imitation performance of the whole imitation process (0-6s) but not the speed, while late phase FEIs selectively increased the speed of successful imitation behavior but not the whole process (other than for the first 2s).

### Spatiotemporal dynamic of MNS representations during repetitive facial expression imitation

We aimed to explore the spatiotemporal dynamic of MNS representations during three fNIRS sessions (2^nd^, 9^th^ and 16^th^ FEI sessions) with neural representations in the MNS and its subregions (i.e. bilateral IFG, STS and IPL), respectively. Considering the classical hemodynamic delay of 6s, the neural pattern similarity (NPS) was assessed for two emotion valence (within positive emotion and within negative emotions) at each time point of imitation via adjusted cosine similarity (see ***Methods***), and the AUC of the NPS curve across imitation sessions was also calculated. We first conducted a set of 2 (emotion valence) × 3 (fNIRS sessions) repeated measures ANOVAs for AUC values for each of the NPS regions. The two-way interaction effect between emotion valence and fNIRS sessions was significant across the overall MNS and for the IFG and STS (*F*(2, 158) > 5.11, *p* < 0.008, *ƞ^2^_p_* > 0.060) but not in IPL (*F*(2, 158) = 2.489, *p* = 0.086, *ƞ^2^_p_* = 0.031). Main effects of emotion valence were observed in the MNS and all its subregions (*F*(1, 79) > 12.78, *p* < 0.002, *ƞ^2^_p_* ≥ 0.139), with greater AUC values for NPS within positive than negative emotions (**Figure 3A**), revealing that the MNS and its subregions have more similar neural representation during the imitation of positive emotion.

**Figure 3.**
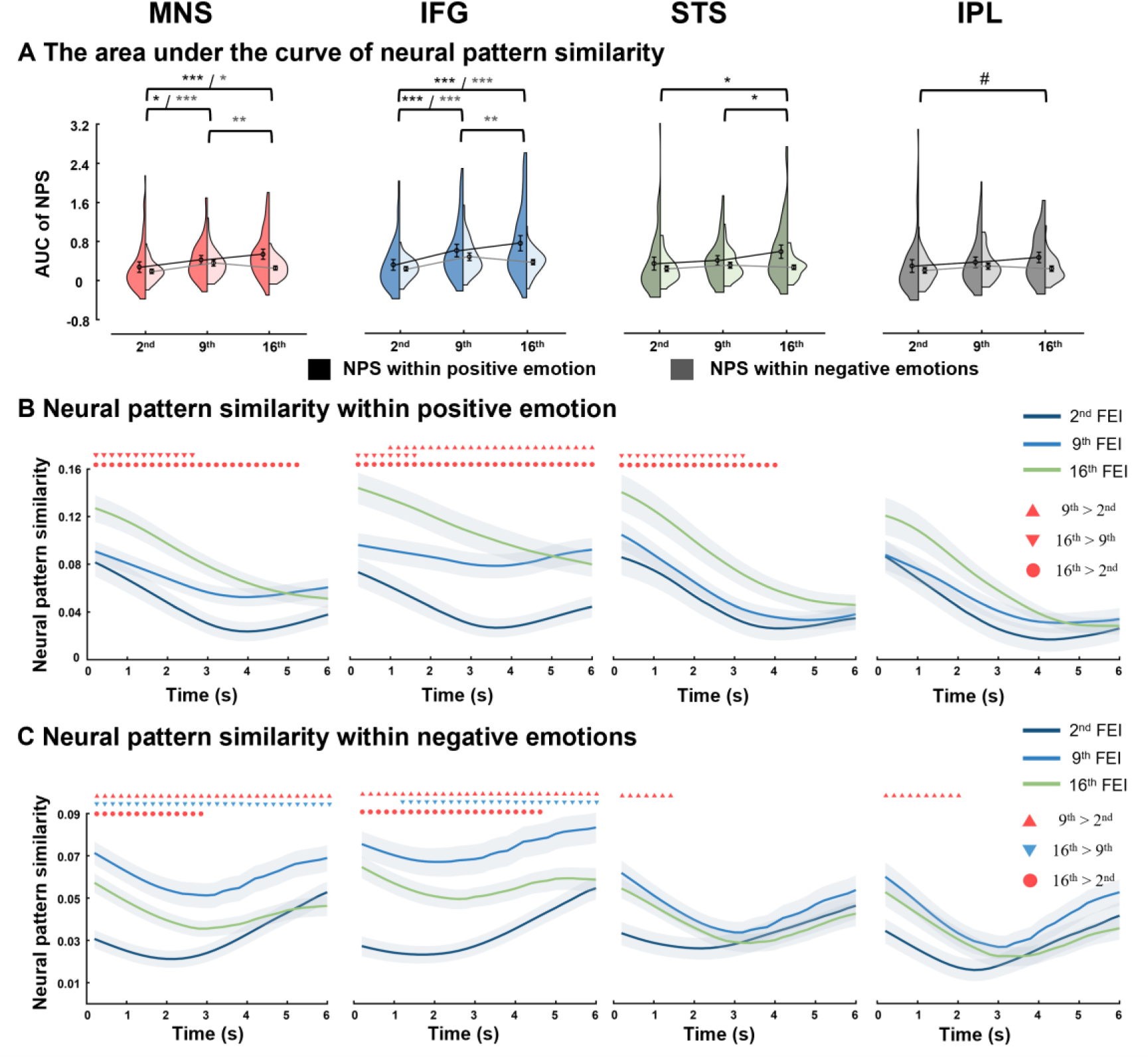
The neural pattern similarity of the MNS and its subregions in three fNIRS sessions. (A) The area under the curve (AUC) of neural pattern similarity for MNS and its subregions for three fNIRS sessions. Data are visualized as split-violin plots with black circles indicating the means and error bars showing the standard errors. * *p* < 0.05, ** *p* < 0.01, *** *p* < 0.001, p with FDR-corrected. (B and C) Time series analyses for neural pattern similarity within imitation of positive and negative emotions. Red upper triangles indicate that the neural pattern similarity (NPS) of the 9^th^ FEI was significantly greater than that of the 2^nd^ FEI using paired sample t-test. Red lower triangles indicate that the NPS of the 16^th^ FEI was greater than the 9^th^ FEI. Blue lower triangles indicate that the NPS of the 16^th^ FEI was smaller than the 9^th^ FEI. Red dots indicate that the NPS of 16^th^ FEI was greater than the 2^nd^ FEI. Error bands represent standard errors. All *p* values were FDR-corrected. AUC = area under the curve; NPS = neural pattern similarity; MNS = mirror neuron system; IFG = inferior frontal gyrus; STS = superior temporal sulcus; IPL = inferior parietal lobule.

Further paired t-test analyses (2^nd^ vs. 9^th^ vs. 16^th^) showed how the neural representation of the MNS dynamic changed. For the long-term effects of imitation (16^th^ FEI > 2^nd^ FEI), the AUC values for positive emotion were significant greater in the 16^th^ FEI than in the 2^nd^ FEI session in the overall MNS, IFG and STS (*t*(56) > 2.92, *p_FDR_* < 0.014) and IPL(*t*(56) = 2.11, *p* = 0.038, uncorrected). For the 9^th^ FEI vs. 2^nd^ FEI, the enhancement effect was also observed in the overall MNS (*t*(56) = 2.17, *p_FDR_* = 0.049) and IFG (*t*(56) = 3.59, *p_FDR_* = 0.001), but in the STS only for the 16^th^ FEI vs. 9^th^ FEI (*t*(56) = 2.42, *p_FDR_* = 0.027). For the NPS within negative emotions, the AUC values were significantly increased in the overall MNS and IFG for the 9^th^ FEI vs. 2^nd^ FEI (MNS: *t*(56) = 4.78, IFG: *t*(56) = 5.78, *ps* < 0.001) and for the 16^th^ FEI vs. 2^nd^ FEI (MNS: *t*(56) = 2.40, *p_FDR_* = 0.019; IFG: *t*(56) = 4.07, *p_FDR_* = 0.001). On the contrary, AUC values (16^th^ FEI vs. 9^th^ FEI) were decreased in the overall MNS (*t*(56) = −3.30, *p_FDR_* = 0.002) and IFG (*t*(56) = −2.97, *p_FDR_* = 0.004).

Subsequently, time series analyses were also conducted to explore the spatiotemporal difference of neural representation among the overall MNS and its subregions (**Figure 3B and 3C**). In contrast to our behavioral finding, the NPS results suggested that greater imitation performance was followed by decreased NPS as the time spent imitating increased. The NPS values for positive emotion (**Figure 3B**) were significant greater in 16^th^ FEI than in 2^nd^ FEI session for the overall MNS (0-5.2s, *p_FDR_* ≤ 0.035), IFG (0-6s, *p_FDR_* ≤ 0.005) and STS (0-4s, *p_FDR_* ≤ 0.039), but not for the IPL (*p_FDR_* ≥ 0.064). This effect was only observed in IFG (1-6s, *p_FDR_* ≤ 0.046) when comparing the 9^th^ vs. 2^nd^ FEI, however the effect improved in the overall MNS (0-2.6s, *p_FDR_* ≤ 0.041), IFG (0-1.6s, *p_FDR_* ≤ 0.043) and STS (0-3.2s, *p_FDR_* ≤ 0.041) for the 16^th^ vs. 9^th^ FEI. In terms of the NPS within negative emotions (**Figure 3C**), the long-term effects (16^th^ vs. 2^nd^ FEI) showed NPS values increased in the overall MNS (0-2.8s, *p_FDR_* < 0.049) and IFG (0-4.6s, *p_FDR_* < 0.022). In addition, the NPS values were greater in the 9^th^ than the 2^nd^ FEI in the overall MNS (0-6s, *p_FDR_* < 0.023), IFG (0-6s, *p_FDR_* < 0.002), STS (0-1.4s, *p_FDR_* < 0.031) and IPL (0-2s, *p_FDR_* < 0.047). However, the NPS values decreased in the 16^th^ relative to the 9^th^ FEI in the overall MNS (0-2.8s, *p_FDR_* < 0.049) and IFG (1.2-6s, *p_FDR_* < 0.047).

Finally, we calculated the correlation between the AUC for imitation performance and neural similarity across emotion valences (see ***Methods***). The results indicated a significant positive correlation between behavioral performance changes and NPS changes when imitating positive emotion in the overall MNS (*r*(79) = 0.278, *p_FDR_* = 0.025, **Figure S1**) and IFG (*r*(79) = 0.337, *p_FDR_* = 0.009, **Figure S2**), but not in the STS (*r*(79) = 0.181, *p* = 0.108) or IPL (*r*(79) = 0.131, *p* = 0.245) or during imitation of negative emotions (*p*s > 0.228). Overall, the analyses indicated that the consistency of MNS representation (particularly in IFG and STS) was improved and predicted imitation performance when imitating positive emotion.

### Representational pathway among MNS subregions

The above analyses revealed the dynamics of the neural representation in the MNS and its subregions during repetitive FEI. However, it was unclear whether the representational pathway within MNS subregions exhibited temporal dynamics. Using directed phase transfer entropy (dPTE, see ***Methods***), we calculated all feasible effectivity connectivities between the NPS time series within the three MNS subregions (from IFG to STS; from STS to IFG; from IFG to IPL; from IPL to IFG; from STS to IPL; from IPL to STS). In addition, the main coupling direction (mCD) of the representational information flow was determined if there was a statistical difference in the dPTE strengths between two regions, e.g., if the dPTE strength from IFG to STS was significantly greater than that from STS to IFG, the mCD between IFG and STS was considered as from IFG to STS. **Figure 4** depicted the inter-regional coupling direction during imitation of positive or negative emotions among the three MNS subregions in the three fNIRS sessions. At the 16^th^ FEI, the mCD of the representational pathway was from STS to IFG (*t*(79) = 2.43, *p*_FDR_ = 0.026, **Figure 4A right**) and from IPL to IFG (t(79) = 3.98, *p*_FDR_ < 0.001) during imitation of positive emotion. For negative emotions, mCD was only observed from IPL to IFG (*t*(79) = 3.79, *p*_FDR_ < 0.001, **Figure 4B right**). No mCD was observed among the three regions in the 2^nd^ and 9^th^ FEIs (*p*s > 0.05, **Figure 4A left** and **4B left**). Furthermore, there were no significant differences in these dPTEs between imitating positive and negative emotions in all three fNIRS sessions (*p*s > 0.05). Detailed statistical results on dPTE values are shown in the **Figure S3**. These results revealed that the flow of information gradually moves into the direction of the IFG following as the number of repetitive FEIs increases.

**Figure 4.**
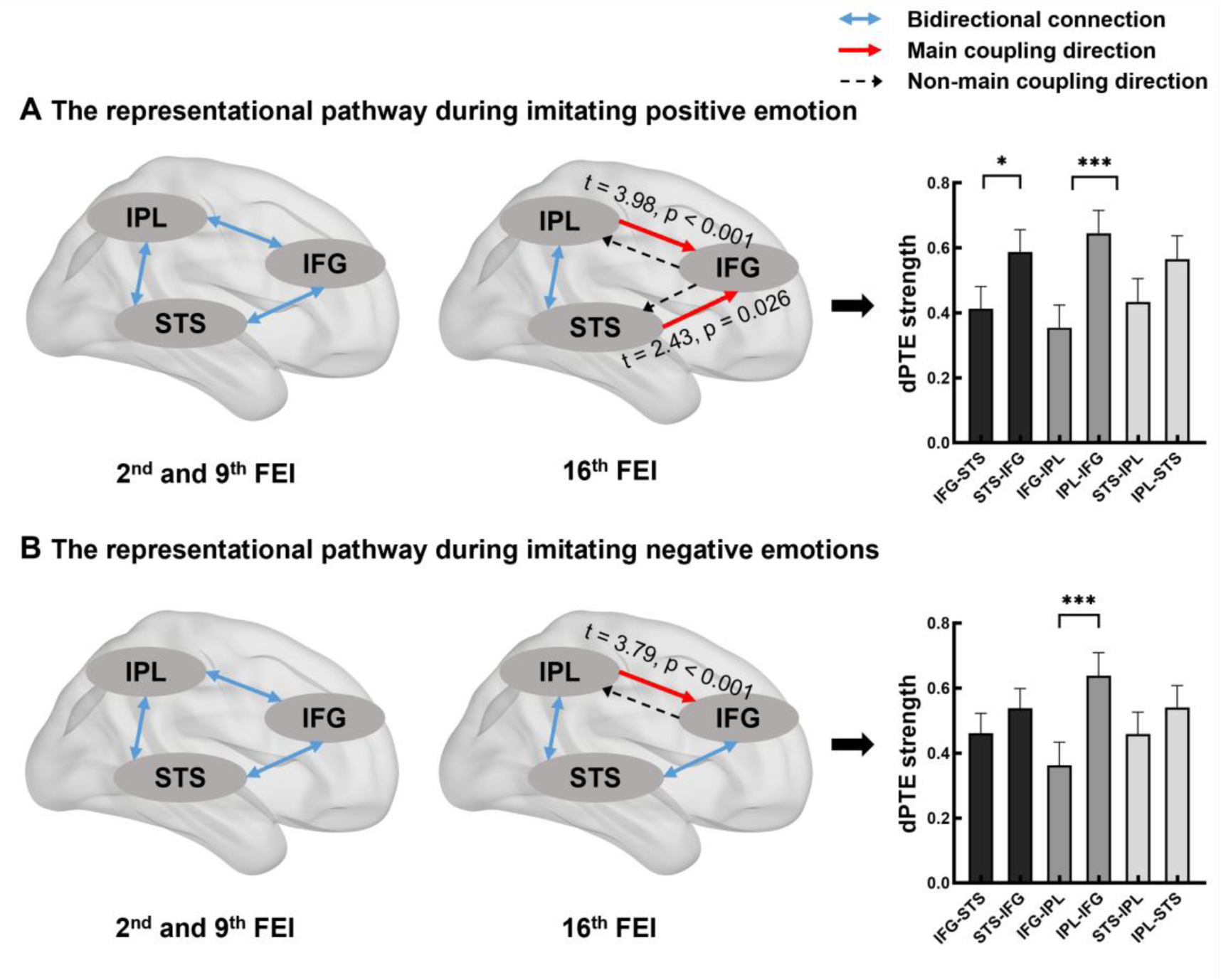
The main coupling directions (mCDs) among MNS subregions during imitation of positive (A) and negative emotions (B). Left: mCDs of the 2^nd^ and 9^th^ FEI; Middle: mCDs of the16^th^ FEI; Right: the dPTE strength among MNS subregions of the 16^th^ FEI. Error bars represent 95% confidence intervals. * *p* < 0.05, \*\*\**p* < 0.001, FDR-corrected. MNS = mirror neuron system; IFG = inferior frontal gyrus; STS = superior temporal sulcus; IPL = inferior parietal lobule; dPTE: directed phase transfer entropy; FEI: facial expression imitation.

### MNS representations predicts facial expression perception

We next systematically validated whether the neural representations of the MNS and its subregions altered neural processing of emotion perception (i.e. transfer effect) after repetitive FEIs. Two pre- and post-FEI fMRI tasks (observe emotional images of faces and scenes vs. neutral images) were collected (see ***Methods***). Specifically, linear support vector machines (SVM, leave-one-subject-out-cross-validation) were trained to determine the accuracy of predicting emotional images (positive/negative) from neutral images via multivariate pattern analyses (MVPA, see ***Methods***) for emotional face and emotional scene tasks respectively, and then we tested whether the classification accuracy in post-FEI was higher than in pre-FEI. For the face task, both pre-FEI and post-FEI sessions were able to significantly classify positive and neutral faces (*p*s < 0.01, **Figure 5A**). Moreover, 10,000 times permutation tests showed that the accuracy of using the IFG as a restriction region for predicting positive and neutral expression faces was significantly higher in the post-FEI than in the pre-FEI session (increase in AUC: 0.134, permutation test *p* = 0.047). Similarly, after the whole repetitive FEI, the accuracy of predicting negative compared to neutral expression faces was improved to varying degrees, regardless of any ROIs used as restriction regions (increase in AUC: 0.042 - 0.128, **Figure 5B**). However, repetitive FEI did not improve the accuracy of predicting emotional scene pictures (permutation test *ps* > 0.05, **Figure 5C and 5D**). These results suggested that the transfer effects of the repetitive FEIs appeared for face emotions (only specific to positive ones) but not for emotional scenes.

**Figure 5.**
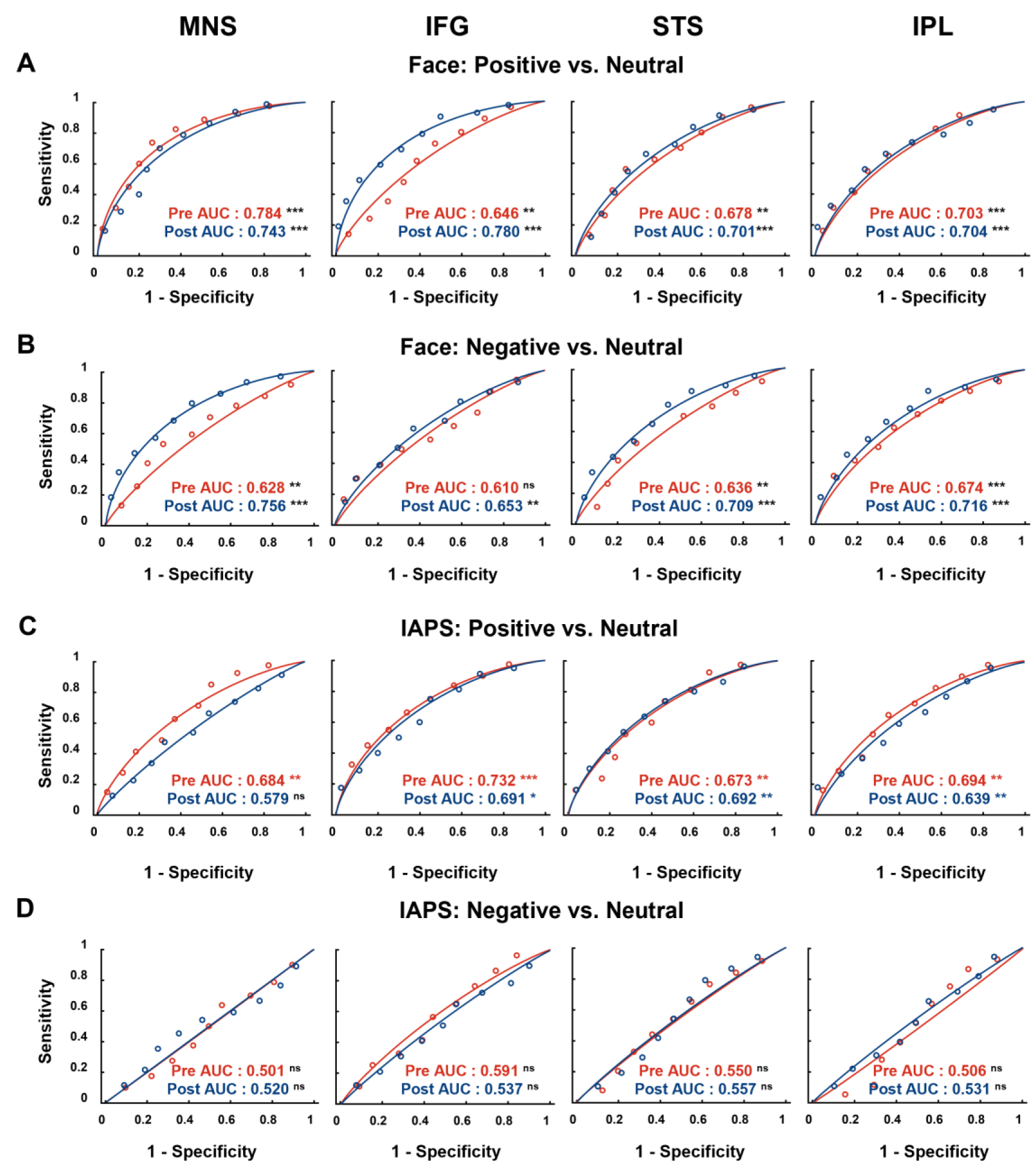
Multivariate pattern analyses with the representations of MNS and its subregions as predictive variables in face emotion expression (A and B) and IAPS (C and D) tasks. Pre/Post AUC = area under the curve of pre/post-FEI; MNS = mirror neuron system; IFG = inferior frontal gyrus; STS = superior temporal sulcus; IPL = inferior parietal lobule; IAPS = International Affective Picture System; * *p* < 0.05, ** *p* < 0.01, \*\*\**p* < 0.001, ^ns^ *p* > 0.05.

### MNS informational connectivity analyses

The above analyses suggested the critical involvement of the MNS, and especially the IFG, in the process of positive facial expression imitation and perception. Given the possible involvement of cortical and subcortical representations during face emotion perception, we used a multi-voxel method known as informational connectivity (IC, see **Figure 6A** and ***Methods***) to explore the representational functional connection between MNS/IFG and other cortical or subcortical regions in discriminating between the activation patterns of positive and neutral emotion face images. Using the MNS as the encompassed region, the IC between MNS and left orbital frontal cortex (LOFC), right orbital gyrus (ROrG), left precentral gyrus (LPrG), right superior temporal gyrus (RSTG), left precuneus, right insula, right inferior temporal gyrus (RITG), right parahippocampal gyrus (RPhG), left precuneus, left cingulate gyrus, left medioventral occipital cortex (LMVOcC), left amygdala and basal ganglia increased after the whole period of repetitive FEIs (**Figure 6B left**, *Z* > 1.96, permutation *p* < 0.05, two-tailed, see **Table S2** for details). With the IFG as the seed region, the IC between orbital gyrus (OrG), STG, fusiform gyrus (FuG), left superior parietal lobule (LSPL), right cingulate gyrus, MVOcC, basal ganglia and right thalamus increased after the whole period of repetitive FEIs (**Figure 6B right**, *Z* > 1.96, permutation *p* < 0.05, two-tailed, see **Table S3** for details).

**Figure 6.**
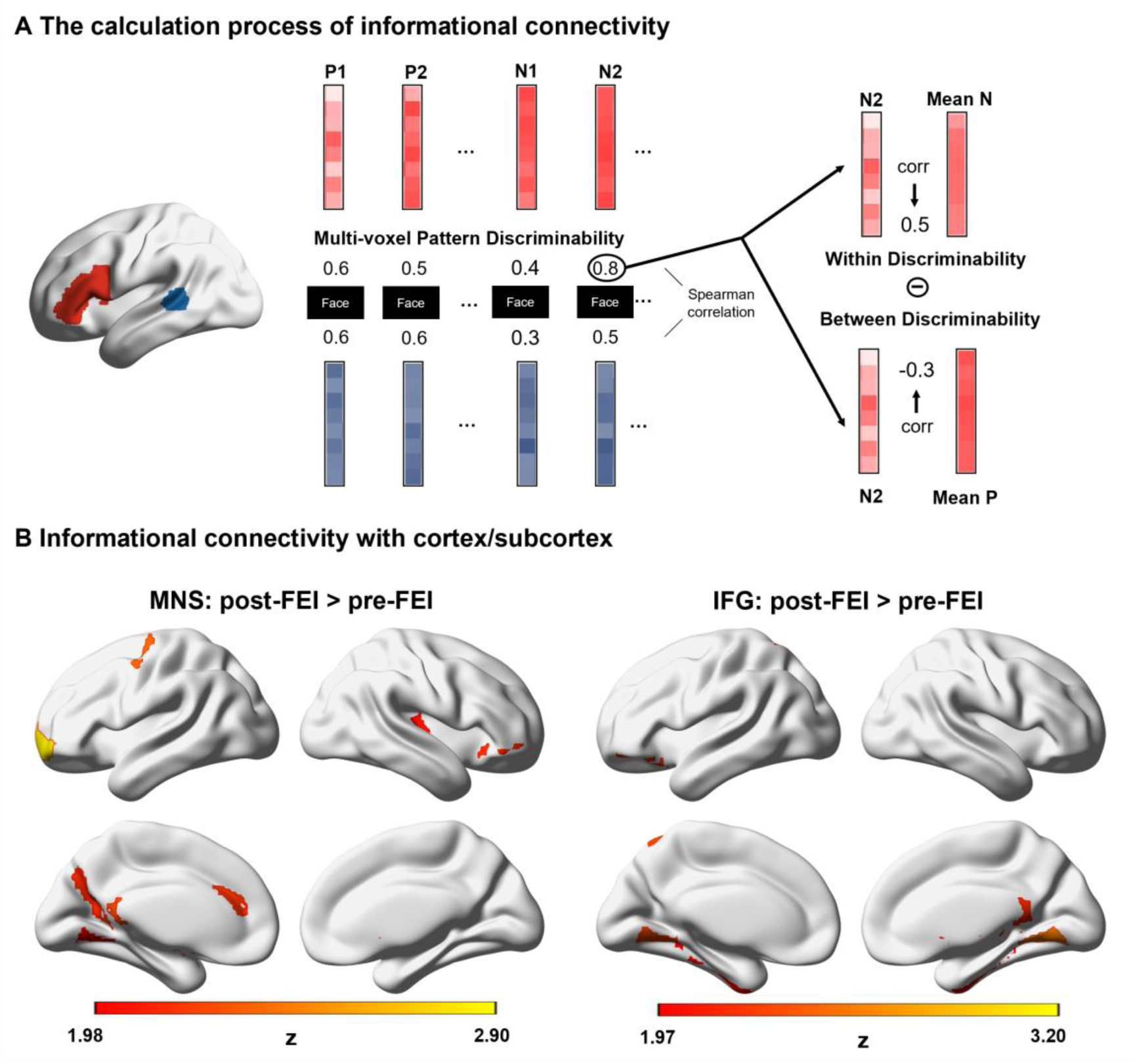
Schematic depiction of informational connectivity analysis methods and results. (A) Informational connectivity between two regions is calculated by obtaining the multi-voxel pattern discrimination (positive vs. neutral emotion) for each trial and region for a given subject and correlating these time series pairwise. (B) Informational connectivity between the MNS/IFG and cortex/subcortex. P1/N1 = the activation pattern of positive/neutral emotion for each trial; Mean P/N = the mean activation pattern across all trials of positive/neutral emotion; corr = Pearson correlation; FEI: facial expression imitation; MNS = mirror neuron system; IFG = inferior frontal gyrus.

### Mediation analysis

To further determine the underlying neural mechanism for the improved positive imitation performance following FEI, a parallel mediation analysis was conducted via R (version 4.2.3) and indicated that the NPS (fNIRS data) of the MNS and the IC (fMRI data) between the MNS and OFC co-mediated the improvement in positive emotion imitation performance (**Figure 7**). The results revealed that the total and direct effect of repetitive FEI on imitation performance of positive emotion were both significant (path *c* = 2.265, *p* = 0.018; path *c*’ = 2.010, *p* = 0.037). Specifically, the improvement in positive imitation performance by repetitive FEI was mediated by NPS of the MNS (indirect effect 1 = 0.834, 95% CI = [0.312, 1.443]) and the IC between MNS and OFC (indirect effect 2 = −0.580, 95% CI = [−1.281, −0.098). A further comparison regarding the size of the mediation effect (Indirect 1 - Indirect 2 =1.414, 95% CI = [0.632, 2.349], bootstrap = 5000) showed that the increased NPS of the MNS and the enhanced IC between the MNS and OFC equivalently mediated the improvement in positive imitation performance by repetitive FEI (**Figure 7**).

**Figure 7.**
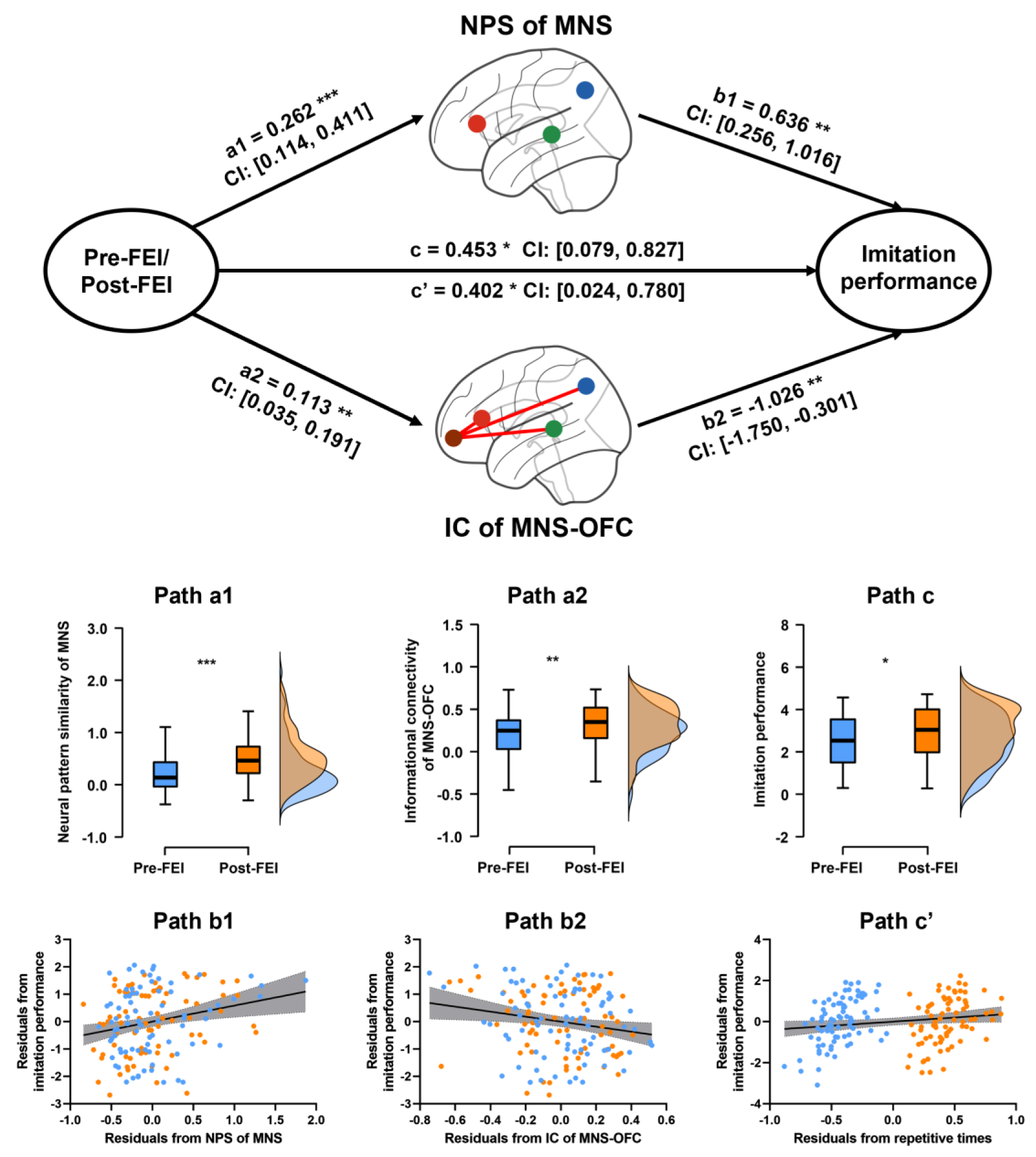
Parallel mediation analysis and examples of each path in the mediation path diagram. FEI: facial expression imitation; MNS = mirror neuron system; OFC = orbital frontal cortex. * *p* < 0.05, ** *p* < 0.01, *** *p* < 0.001.

## Discussion

Since the discovery of MNS in humans two decades ago^9,39,40^, the essence of the mirror mechanism^2,12,41^ and its translational relevance in a variety of clinical conditions^10,16^ has been explored. Notably, among these studies, motor imitation has been the primary focus^21,42,43^. During dynamic social interaction, the ability to recognize facial expressions or mirroring other’s emotions may represent a basic prerequisite for the development of social cognition^44^. However, the impact of repetitive imitation or mirroring of facial expressions on imitation performance and processing in the MNS remains unclear. Thus, in the current study, we investigated the effects of repetitive imitation (16 times during one month) of facial expressions (positive vs. negative) in a naturalistic environment and fNIRS data collected at three different time points were used to explore how spatiotemporal encoding within the MNS changed across the period of repeated FEIs. More importantly, alterations in the patterns of MNS activity and its connectivity with other cortical and subcortical regions during emotion perception was also investigated at pre- and post-repetitive imitation times points using fMRI. Overall, the current findings suggest that behavioral imitation performance, neural representations and the information flow within the MNS exhibit valence-specific improvement following the repetition of imitation and are paralleled by enhanced patterns predictive of which face emotions are being perceived.

Our findings are compatible with one previous study showing evidence for valence-specific imitation patterns with frontal regions involved in imitation of negative emotions (i.e. angry and sad) and occipitotemporal and subcortical regions for positive ones (i.e. happy) was found^45^ and another study reporting a positive emotion advantage effect in social attention^46^. Together they suggest that there may be a difference between imitation of positive and negative emotions. Additionally, within the MNS the similarity pattern of imitation varies between positive and negative face emotions (angry, sad and fear)^24^. Critically, the results indicate that better imitation performance and greater similarity within the MNS occurred after repeated imitation of positive relative to negative face emotions (i.e. angry, sad, fear and disgust) at both group and individual levels. This may be due to positive emotion imitation promoting increased attention^46^ and that greater accuracy indicates increased importance for social interaction^47^.

Given the high temporal resolution of fNIRS imaging, we can explore the spatiotemporal dynamic neural representation of MNS at different time points (2^nd^, 9^th^, 16^th^ FEI) during the whole period of repeated FEIs. The results suggested that the MNS pattern similarity gradually increased for positive emotion (16^th^ > 9^th^ > 2^nd^ FEI), indicating that this improvement in the neural pattern similarity may be advantageous for performance^48,49^. By contrast, one fMRI study in which participants were required to observe repeated movies with the same movement and the activity, both the IPL and IFG exhibited suppression^50^. This may due to the requirement for participants to only perceive repeated movements without having to actually imitate them whereas in our study participants were specifically instructed to imitate observed face emotions as vividly as possible and informed that their performance would be assessed. One issue that should be noted is that greater neural pattern similarity was found at the middle compared to the late phase (9^th^ >16^th^ FEI) of negative emotion imitation, which may indicate initial unconscious avoidance behaviors towards negative or threatening stimuli^47^, although the subjects were required to consciously imitate the expressions as vividly as possible. Specifically, the representational function of MNS subregions exhibited divergent roles with the IFG progressively engaged from the early to late phases of the whole repetitive imitation process, while the STS was only involved in the late phase (from 9^th^ to 16^th^ FEI) and the IPL not showing any time-dependent changes. Taken together findings suggest that the whole MNS may be involved during the early phase of repetition with the IFG becoming progressively more involved over time. As such, the IFG seems to be particularly engaged in the whole process of imitation including understanding actions as well as copying and executing them. On the contrary, the STS as a major region involved in for visual processing and interpretation of stimuli may be slower to develop imitation dependent changes reflecting a greater cognitive role. The IPL contributed to a similar degree across the three time points, possibly indicating that it contributes less to improved MNS processing with repeated FEIs over time.

Additionally, the detailed time-series results for neural pattern similarity also supported the above findings for a divergent role of MNS subregions during repetitive FEIs. During the last repetition of imitation (i.e. the 16^th^ FEI) MNS neural pattern similarity was greater than during the first one (2^nd^ FEI) for happy expressions across all 30 of the individual time frames during the 6s period of imitation, although when contrasting the middle (9^th^) and late (16s) phases this was only the case for the time frames during the first 3 s. In addition, the dynamic changes in IFG neural pattern similarity showed differences with those of the STS and IPL. In line with the neural pattern similarity results the IPL did not show any differences in individual time frames across the three repetition phases. On the one hand, the IFG exhibited differences between the last FEI time point and the other two earlier ones and using a short time window of ∼2s there was also a difference between 9^th^ and 16^th^ FEI time points. On the other hand, for the STS pattern similarity there was only improvement using a short time period of ∼3s) when comparing the late (16^th^) with the early (2^nd^) or middle (9^th^) time-points. Together with the neural pattern similarity results, these findings suggest that whereas pattern similarity in the IFG may involve the total 6 s imitation time window for the STS it may only involve the first 3 s and for the IPL there was no indication of time-frame dependent changes across repetitions. A significant positive correlation between the improvement in MNS/IFG neural pattern similarity and enhanced positive expression imitation performance was found suggesting that repetition of positive emotion imitation is most closely linked with neural representation in the MNS and particularly the IFG. However, this appears to be less the case for repetitive negative emotion imitation.

Furthermore, previous studies have proposed that the mechanism of imitation was driven by the bidirectional connections between the IFG and IPL, and the IPL and STS^10,41^. In line with this, we confirmed that during the early and middle phases of repeated emotion imitation, IFG-IPL, IPL-STS and STS-IFG all exhibited bidirectional connections and they contributed equally to emotion imitation performance. However, by the end of the period of repetitive emotion imitation the main flow of representational information was directed from the IPL/STS to the IFG. Thus, after a period of repeated imitation the IFG may play a more dominant role in MNS processing, receiving increased information about sensory or visual inputs in order to copy or understand the goal (i.e. to imitate behaviors with greater accuracy and/or faster). This may reflect the important role that the IFG has in emotion recognition and emotion evaluation^51^ as well as in imitation learning^52^.

We found that long-term repetitive imitation enhanced neural responses to perception of positive face emotion but not positive emotional social scenes. Additionally, repetitive imitation of face emotions increased the representational connectivity between the IFG in the MNS and other frontal and subcortical regions involved in recognition and affective processing of faces (notably the fusiform gyrus, medial frontal gyrus and amygdala) and reward (orbitofrontal cortex)^53,54^ suggesting that it could be a potential intervention for individuals with problems in recognition and responses to face emotion expressions as well as evaluation of their rewarding properties. Importantly, the greater MNS neural pattern similarity observed in fNIRS data and increased MNS-OFC informational connectivity derived from fMRI data contributed equally to improving positive imitation performance. Furthermore, the greater influence of repetitive FEI on the processing and reward value of positive emotion faces may indicate that it might have therapeutic value in individuals suffering from mood as well as neurodevelopmental disorders associated with reduced responsivity to positive emotion.

The present study has the following limitations. Firstly, our fNIRS recordings were only made from the three main cortical MNS regions and did not include some other motor regions (i.e. premotor and supplementary motor regions), although previous studies have suggested that STS and IPL also represent motoric description of the action ^10^. Secondly, we only collected data from male subjects for the current study and therefore we cannot extrapolate our findings to females. Thirdly while our imitation protocols focused on deliberate conscious imitation of face expressions we cannot completely rule out some contributions from unconscious motor contagion effects, particularly for smiling^55,56^, although evidence suggests that neural processing of emotional contagion may differ somewhat to that for conscious imitation^57^.

In summary, our results suggest that repeated imitation of face emotions improves the accuracy of imitation behaviors and influences spatiotemporal dynamics of MNS representations with divergent effects on the three main cortical MNS regions. Specifically, the IFG plays the role of integrating visual inputs and contributes to both more accurate and rapid imitation. In addition, long-term repetitive imitation promoted alterations in MNS informational connectivity with other cortical and subcortical regions subserving processing and reward components of face and face emotion processing during perception of positive emotion faces. The current results therefore provide a more detailed understanding of how repetitive imitation shapes spatiotemporal processing in the MNS to both facilitate imitation accuracy as well as processing of faces and face emotions. As such they pave the way for a variety of potential clinical and translational applications which utilize emotion imitation as a strategy to improve both neural and behavioral control of social cognition and motivation.

## Methods

### Participants

One hundred healthy right-handed male (based on their self-reports) participants were recruited from the University of Electronic Science and Technology of China via the campus bulletin board system (BBS). All participants self-reported having normal or corrected-to-normal vision, being free of any medical or psychiatric disorders and not taking current or regular medication. Twenty participants were excluded due to technical problems with FaceReader recording (n = 2), incomplete fNIRS data for the three recording sessions (n = 9) and incomplete fMRI data for two scans (n = 9), resulting in 80 participants (mean age = 21.19 ± 2.14 years) included for further analyses. All participants were required to complete facial expression imitation (FEI) on 16 different occasions with three fixed fNIRS sessions (during 2^nd^, 9^th^ and 16^th^ FEI) over a period of averaged one month (3-4 times of FEI per week). They were also required to complete fMRI scans before and after the repetitive FEIs where they performed two task paradigms (face emotion perception and emotional scene perception) (**Figure 1A**). All participants provided written informed consent, and this study was approved by the Institutional Review Board of the University of Electronic Science and Technology of China and conformed with the latest revision of the declaration of Helsinki.

### Experimental Protocol

#### Repetitive FEIs Procedure

For each FEI session, 60 validated facial expressions ^24,58^ with positive (i.e. happy) and negative (i.e. angry, disgust, fearful and sad) emotions (4 ∗ 5 expression * 3 run) were presented randomly on a 27-inch monitor at a resolution of 1920 × 1080 pixels (60 Hz). The stimuli were matched and not repeated across the 16 FEI sessions. Participants were required to sit 60 cm away from the presentation monitor and asked to imitate the presented facial expressions as accurately as possible over a period of 6s ^59^. A blank screen was displayed between each imitation trial for 3s as a fixed inter-trial interval (**Figure 1B**). Between runs, a break of 30 to 60 seconds was taken to prevent possible fatigue. In addition, imitation performance was recorded and measured via automatic facial expression analysis software - FaceReader (Noldus 7.1) and fNIRS was used to detect activity changes in the MNS (**Figure 1C**, see ‘**fNIRS data collection and analyses** for details) during the 2^nd^, 9^th^ and 16^th^ FEI sessions.

#### fMRI Procedures

Before and after (1-3 days) the period of repetitive FEIs (Pre-FEI and Post-FEI sessions), participants were asked to complete emotional face perception and emotional scene perception (International Affective Picture System-IAPS) tasks (**Figure 1D**) in the fMRI scanner. The order of the two tasks was counterbalanced across participants. Both emotional perception tasks were presented via E-prime 2.0 (Psychology Software Tools), and each task comprised two runs. Each run commenced with a 10 s and ended with a 4 s blank screen to achieve a stable signal. For the event-designed face perception task, 64 validated facial expressions and 16 scrambled pictures (8 × 5 categories [positive: happy; negative: angry, fearful; neutral and scramble] × 2 runs) were presented randomly. In each trial, a face picture was displayed for 4s and participants were instructed to perceive the emotion expressed by the person, followed by a white fixation jitter for 2-4s (0.2s gap). For the block-designed IAPS perception task, 72 emotional scene pictures and 24 scrambled pictures (4 × 6 blocks × 4 categories [positive, negative, neutral and scramble] × 2 runs) were presented randomly. In each block, four scene pictures were continually displayed for 14s (each one for 3s with a 0.5 s fixation) and participants instructed to perceive the emotions being displayed by the individuals in the scenes. The inter-block interval was 10-14s (2s gap). For the two fMRI scans, identical task paradigms with different and matched stimulus materials were employed, and the pictures of emotional faces were different from those in the repetitive FEI phase. An independent sample of male participants (n=31, Mean age = 21.23 ± 2.04 years) rated the intensity (from 1-9 points using visual analog scale) in terms of negative and positive face stimuli and IAPS stimuli and no significant differences were found between the two tasks across valences (positive: *t*(30) =0.925, *p* =0.362; negative: *t*(30) =0.687, *p* =0.498).

#### Behavioral data collection and analysis

During each FEI session, an experimenter mounted a video camera (Logitech HD Pro-Webcam C920, Logitech, Switzerland) on the presentation monitor and adjusted the camera so that the video frame only included the full face of the participant. Imitation performance values ranged from 0 to 1 for each of five facial expression elements (happy, angry, disgust, fearful and sad) and were automatically evaluated separately by FaceReader in 15Hz, with higher values indicating higher accuracy of facial expression imitation. Next, we down-sampled the imitation performance to 5.09Hz by the “*resample*” function in MATLAB 2019b to keep consistent with the fNIRS data. Due to the potential challenges of unfamiliarity with imitation and the negative impact of wearing the fNIRS electrode cap on imitation accuracy, only data for imitation performance from 3^rd^-8^th^ (early-repetitive FEI phase) and 10-15^th^ (late-repetitive FEI phase) FEI sessions were used for further analyses.

The area under the curve (AUC) of imitation performance across each individual imitation (6s) was calculated for imitating positive expression (happy) and negative expressions (average of angry, disgust, fearful and sad), respectively, with the AUC chosen for statistical analyses because of its superior sensitivity ^60^. The formula for calculating the AUC is represented as follows.

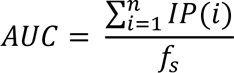

Where, *n* represents the number of onset-times for the imitation process, IP denotes the imitation performance, and *f*_s_ denotes the sampling frequency of imitation performance (5Hz).

Next, a 2 by 12 mixed-effects ANOVA was conducted to explore the main effects and interaction between emotion valence (i.e., positive and negative) and the number of repetitions (from 3^rd^-8^th^ and 10-15^th^ FEI). Furthermore, repeated measures correlation ^61^ was employed to examine whether the AUC of imitation performance and imitation performance at each time point (during the 6s of imitation process) for positive and negative expressions improved (or decreased) significantly with the number of repetitions (across whole-early- and late-repetitive FEI phases). In addition, we defined the onset-time at which imitation performance improves to 90% of the difference between the best imitation performance and the imitation performance at 0s as the onset-time of successful imitation. Similarly, a 2 by 12 mixed-effects ANOVA and repeated measures correlations were conducted to confirm the effects of emotion valence and number of repetitions on the onset-time for successful imitation. ANOVAs and repeated measures correlations were conducted via SPSS 25.0 (IBM SPSS Statistics) and R (version 4.2.3), respectively.

### fNIRS data collection and analyses

#### fNIRS acquisition and preprocessing

During the 2^nd^, 9^th^ and 16^th^ FEI sessions, a NIRSport2 system (NIRx Medical Technologies LLC, Berlin, Germany) with LED light sources (sampling rate: 5.09Hz, wavelengths: 760nm and 850nm) was used to collect hemodynamic data from the MNS. Two identical optode probe sets of 8 sources and 8 detectors with a maximum inter-optode distance of 3 cm were used based on fOLD v2.2 ^62^, resulting in 42 channels (21 channels in each brain hemisphere). The optode probes covered the three key MNS regions, including IFG, IPL and STS and were arranged on the NIRS cap according to the international 10-10 system (**Figure 1C**) based on previous studies ^63–65^. The Montreal Neurological Institute (MNI) coordinates of the probes and information on generated channels were provided in **Table S4**. The fNIRS data preprocessing was carried out using the NIRS-KIT toolbox ^66^ in MATLAB 2019b. Specifically, the modified Beer-Lambert law transformed raw data into oxyhemoglobin (HbO) and deoxyhemoglobin (HbR) concentrations. Next, a polynomial regression model was applied to estimate a linear trend, which was then subtracted from the raw hemoglobin concentration signal through detrending. To eliminate motion artifacts, the temporal derivative distribution repair (TDDR) algorithm ^67^ was applied to all signals. To effectively filter noise and recover the hemodynamic response without distorting the signal phase ^68^, a 500th order band-pass FIR filter (0.01-0.08 Hz) was employed in the final step. This study focused on changes in HbO concentrations due to its superior signal-to-noise ratio in fNIRS measurements ^69^.

#### Neural pattern similarity

We calculated the neural pattern similarity (NPS) during imitation of positive and negative emotions in the overall MNS and its three individual subregions during the three fNIRS scans via adjusted cosine similarity ^70^, which can remedy the potential effect of cosine similarity and Pearson correlation on numerical insensitivity. The formula for calculating the NPS (range −1 to 1) between two imitation trials (*x*_1_, *x*_2_) at time *t* is represented as follows.

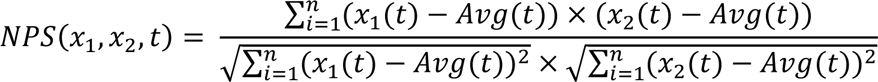

Where, *n* represents the number of channels for the restricted region of interest (ROI: MNS, IFG, IPL and STS), *x*_1_ (t) and *x*_2_(t) are the hemodynamic response in a restricted ROI at imitation time *t* (considering a classical hemodynamic delay of 6s), and *Avg*(t) is the average hemodynamic response at time *t* across all imitation trials for a participant. Next, the average NPS during imitation of positive and negative emotions was calculated for each participant. Similarly, the AUC of NPS across each individual imitation was evaluated for positive and negative emotions, respectively. Furthermore, a set of 2 (emotion valence) by 3 (fNIRS sessions) mixed-effects ANOVAs and further time series analyses were conducted to explore the emotion valence and repetitive phase effects on the NPS.

#### Direction of representational pathway

Phase transfer entropy (PTE) was employed to assess the representational pathway among MNS subregions during FEI. Taking the representational pathway from IFG to STS as an example, the PTE value can be expressed as:

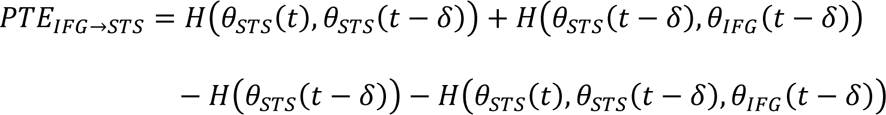

Where *θ*_IFG_(t) and *θ*_STS_(t) are the instantaneous phase of the NPS time series for STS and IFG, respectively, and *δ* is the delay between the source signal and the target signal. Furthermore, the directed PTE (dPTE, range 0 to 1) was used to reduce PTE biases:

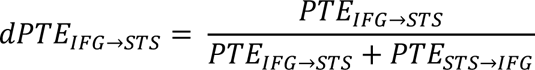

If 0.5 < dPTE_IFG→STS_ ≤ 1, the representational pathway between IFG and STS is considered to be preferentially from IFG to STS. While 0 ≤ dPTE_IFG→STS_ < 0.5, the representational pathway is considered to be preferentially from STS to IFG. In addition, if dPTE_IFG→STS_ is statistically greater than dPTE_STS→IFG_, the main coupling direction (mCD) of the representational information flow was considered as from IFG to STS. We calculated all feasible representational pathways between the NPS time series within the three MNS subregions (from IFG to STS; from STS to IFG; from IFG to IPL; from IPL to IFG; from STS to IPL; from IPL to STS), and then determined the mCD during imitation of positive or negative emotions in the three fNIRS sessions via paired t-tests.

#### fMRI data collection and analyses

The fMRI data acquisition and analyses for the pre-and post-FEI phases were completely consistent.

#### fMRI acquisition and preprocessing

Structural and functional imaging data were acquired on a 3.0T GE Discovery MR750 system (General Electric, Milwaukee, WI, USA) with an 8-channel head-neck coil at the MRI Center at University of Electronic Science and Technology of China. Using a spoiled gradient echo pulse sequence, high-resolution T1-weighted structural images were acquired for the whole brain (repetition time (TR) = 6ms; echo time (TE) = 2ms; number of slices = 156; flip angle (*θ*)= 9^◦^; field of view (FOV) = 256 mm × 256 mm; acquisition matrix = 256 × 256; thickness = 1 mm). Using a T2*-weighted echo planar imaging (EPI) sequence, the high-resolution functional images were acquired (TR = 2000ms; TE = 30ms; *θ* =90^◦^; FOV = 240 mm × 240 mm; acquisition matrix = 64 × 64; slice thickness = 3 mm; and slices = 36 with an interleaved ascending order).

Imaging data were preprocessed using *fMRIPrep* 20.2.1 ^71^, which is based on Nipype 1.5.1 ^72,73^. T1-weighted structural images were corrected for intensity non-uniformity with N4BiasFieldCorrection and were then skull-stripped by antsBrainExtraction.sh, using OASIS30ANTs as a target template. Volume-based spatial normalization to the standard space (MNI152NLin2009cAsym) was performed through nonlinear registration by antsRegistration. For each functional run, the first 5 volumes were discarded to ensure steady-state longitudinal magnetization. A reference volume and its skull-stripped version were generated and the BOLD reference was then co-registered to the corresponding structural data by boundary-based registration with nine degrees of freedom. Functional runs were slice-time corrected using 3dTshift (AFNI) and then resampled to their original space by applying the transforms to correct for head-motion. The BOLD time-series were subsequently normalized to the MNI152NLin2009cAsym space (2 × 2 × 2 mm). The frame-wise displacement (FD) was calculated for each functional run in Nipype. Additionally, the BOLD time-series were filtered temporally with a nonlinear high-pass filter with a 128s cut-off to remove low-frequency drifts. Finally, the functional images were smoothed with a Gaussian kernel at full width at half maximum (FWHM, 8 mm) via Statistical Parametric Mapping (SPM12 v7771, https://www.fil.ion.u cl.ac.uk/spm/software/spm12).

#### Univariate analysis

The General Linear Model (GLM) was conducted on the smoothed-fMRI data using SPM 12 v7771. On the individual level, the face emotion and IAPS tasks were modeled with a double gamma hemodynamic response function (HRF) on four separate regressors for the four emotional conditions (positive, negative, neutral and scrambled emotional stimuli). Confound regressors in the SPM statistical model included six head movement parameters and their squares, their derivatives and squared derivatives, leading to 24 motion-related nuisance regressors in total. Additionally, three contrasts of interest ([positive- scramble], [negative- scramble] and [neutral- scramble]) were produced on the individual level.

#### Multivariate pattern analysis

In order to assess emotion perception (face task and IAPS task) before and after the repetitive FEIs, classifiers were trained to distinguish positive/negative and neutral trials or blocks using multivariate pattern analysis (MVPA, Chang et al., 2015). We employed a support vector machine (SVM, leave-one-subject-out-cross-validation) algorithm with linear kernel (C = 1) implemented in CanlabCore tools (https://github.com/canlab/CanlabCore) with the individual con maps within restriction regions (i.e. MNS, IFG, STS and IPL) as features to determine the accuracy of predicting emotional images (positive/negative) from neutral images. Thus, a total of 2 (fMRI task: face and IAPS) × 2 (fMRI time: pre-FEI and post-FEI) × 4 (region of interest-ROI: MNS, IFG, STS and IPL) × 2 (image valence: positive vs. neutral and negative vs. neutral) SVM models were trained respectively. Restriction regions (IFG, STS and IPL) were identified by the Automated Anatomic Labelling (AAL) atlas which was implemented in the WFU Pick Atlas ^74^, and the MNS was defined as the combination of these three regions (see **Figure S4**). In addition, permutation tests (10,000 permutations) were conducted to investigate whether the classification accuracy in post-FEI were significantly higher than pre-FEI.

#### Single-trial estimation

The GLM was also used to compute the *t* map for each of the 80 trials in the face emotion task (4s per trial) and the 24 blocks in the IAPS task (14s per trial), respectively. Consistent with univariate analysis, the single-trial estimation was modeled with the presentation of each trial as a starting point and convolved with the double gamma HRF. The 24 motion-related nuisance regressors were also used in each single-trial model.

#### Informational connectivity between MNS and cortex/subcortex

To investigate whether the representational information in the MNS and in the cortex/subcortex during emotion perception was increased after repetitive FEIs, we used an informational connectivity (IC) approach ^34–36^. The metric underlying IC quantifies the correlation between the time series of multi-voxel pattern discriminability (condition 1 vs. condition 2, i.e. positive vs. neutral and negative vs. neutral) for each trial (**Figure 6A**). Spearman’s correlation coefficients were used and then transformed into Fisher’s Z scores. Taking the representational information between IFG and hippocampus under the condition of positive vs. neutral face emotion as an example, the IC value was quantified with the following procedure:

1. Within condition discriminability is defined as the Pearson correlation coefficient between (i) the vector of voxel activation values (i.e., its activity pattern) for the single positive trial and (ii) the vector of mean voxel activation values across all single positive trials for a given subject, and then transformed into Fisher’s Z scores.
2. Between condition discriminability is defined as the Pearson correlation coefficient between (i) the vector of voxel activation values (i.e., its activity pattern) for a single negative trial and (ii) the vector of mean voxel activation values across all individual negative trials for a given participant, and then transformed into Fisher’s Z scores.
3. Multi-voxel pattern discriminability = Within condition discriminability − Between condition discriminability.
4. IC between IFG and hippocampus is defined as the Spearman correlation between the time series of multi-voxel pattern discriminability for IFG and hippocampus, and then transformed into Fisher’s Z scores.

For this exploratory analysis, we finally used a relatively loose threshold (*Z* > 1.96 and permutation *p* < 0.05, two-tailed) to identify which specific ICs changed after repetitive FEIs.

## Acknowledgements

This work was supported by the Natural Science Foundation of Sichuan Province [grant number 2022NSFSC1375 – W.Z., 2023NSFSC1185 – X.Z.,], Fundamental Research Funds for the Central Universities, UESTC [grant number ZYGX2020J027 – W.Z.], National Natural Science Foundation of China (NSFC) [grant number 31530032 – K.M.K.] and Key Scientific and Technological projects of Guangdong Province [grant number 2018B030335001 – K.M.K.].

## Authorship Contribution Statement

Q.L. and W.Z. designed the experiment and drafted the manuscript. Q.L., X.Z., S.Z, C.L., Y.Y., C.L., X.S. and W.Z. conducted the experiment, and analyzed the data. B.B. and K.M.K. provided feedback and revised the manuscript.

## References

1. Chartrand, T. L. & Lakin, J. L. The Antecedents and Consequences of Human Behavioral Mimicry. Annu. Rev. Psychol. 64, 285–308 (2013).

2. Heyes, C. Causes and consequences of imitation. Trends in Cognitive Sciences 5, 253–261 (2001).

3. Heyes, C. Imitation. Current Biology 31, R228–R232 (2021).

4. Field, T. Imitation enhances social behavior of children with autism spectrum disorder: A review. Behavioral Development Bulletin 22, 86–93 (2017).

5. Field, T. M., Woodson, R., Greenberg, R. & Cohen, D. Discrimination and Imitation of Facial Expression by Neonates. Science 218, 179–181 (1982).

6. Over, H. The Social Function of Imitation in Development. Annu. Rev. Dev. Psychol. 2, 93–109 (2020).

7. Braadbaart, L., De Grauw, H., Perrett, D. I., Waiter, G. D. & Williams, J. H. G. The shared neural basis of empathy and facial imitation accuracy. NeuroImage 84, 367–375 (2014).

8. Bonini, L., Rotunno, C., Arcuri, E. & Gallese, V. Mirror neurons 30 years later: implications and applications. Trends in Cognitive Sciences 26, 767–781 (2022).

9. Iacoboni, M. et al. Cortical Mechanisms of Human Imitation. Science 286, 2526–2528 (1999).

10. Iacoboni, M. & Dapretto, M. The mirror neuron system and the consequences of its dysfunction. Nat Rev Neurosci 7, 942–951 (2006).

11. Molenberghs, P., Brander, C., Mattingley, J. B. & Cunnington, R. The role of the superior temporal sulcus and the mirror neuron system in imitation. Human Brain Mapping 31, 1316–1326 (2010).

12. Iacoboni, M. Neurobiology of imitation. Current Opinion in Neurobiology 19, 661–665 (2009).

13. Iacoboni, M. Imitation, Empathy, and Mirror Neurons. Annu. Rev. Psychol. 60, 653–670 (2009).

14. Molenberghs, P., Cunnington, R. & Mattingley, J. B. Brain regions with mirror properties: A meta-analysis of 125 human fMRI studies. Neuroscience & Biobehavioral Reviews 36, 341–349 (2012).

15. Park, S., Matthews, N. & Gibson, C. Imitation, Simulation, and Schizophrenia. Schizophrenia Bulletin 34, 698–707 (2007).

16. Perkins, T., Stokes, M., McGillivray, J. & Bittar, R. Mirror neuron dysfunction in autism spectrum disorders. Journal of Clinical Neuroscience 17, 1239–1243 (2010).

17. Binder, E. et al. Lesion evidence for a human mirror neuron system. Cortex 90, 125–137 (2017).

18. Bonini, L. & Ferrari, P. F. Evolution of mirror systems: a simple mechanism for complex cognitive functions. Annals of the New York Academy of Sciences 1225, 166–175 (2011).

19. Izard, C. et al. Emotion Knowledge as a Predictor of Social Behavior and Academic Competence in Children at Risk. Psychol Sci 12, 18–23 (2001).

20. de Waal, F. B. M. & Preston, S. D. Mammalian empathy: behavioural manifestations and neural basis. Nat Rev Neurosci 18, 498–509 (2017).

21. Heiser, M., Iacoboni, M., Maeda, F., Marcus, J. & Mazziotta, J. C. The essential role of Broca’s area in imitation. Eur J of Neuroscience 17, 1123–1128 (2003).

22. Hogeveen, J. et al. Task-dependent and distinct roles of the temporoparietal junction and inferior frontal cortex in the control of imitation. Soc Cogn Affect Neurosci 10, 1003–1009 (2015).

23. Restle, J., Murakami, T. & Ziemann, U. Facilitation of speech repetition accuracy by theta burst stimulation of the left posterior inferior frontal gyrus. Neuropsychologia 50, 2026–2031 (2012).

24. Zhao, W. et al. Differential responses in the mirror neuron system during imitation of individual emotional facial expressions and association with autistic traits. NeuroImage 277, 120263 (2023).

25. Audrain, S. & McAndrews, M. P. Schemas provide a scaffold for neocortical integration of new memories over time. Nat Commun 13, 5795 (2022).

26. Lu, Y., Wang, C., Chen, C. & Xue, G. Spatiotemporal Neural Pattern Similarity Supports Episodic Memory. Current Biology 25, 780–785 (2015).

27. Xue, G. et al. Greater Neural Pattern Similarity Across Repetitions Is Associated with Better Memory. Science 330, 97–101 (2010).

28. Ekhlasi, A., Nasrabadi, A. M. & Mohammadi, M. R. Direction of information flow between brain regions in ADHD and healthy children based on EEG by using directed phase transfer entropy. Cogn Neurodyn 15, 975–986 (2021).

29. Li, M. et al. Transitions in information processing dynamics at the whole-brain network level are driven by alterations in neural gain. PLoS Comput Biol 15, e1006957 (2019).

30. Lobier, M., Siebenhühner, F., Palva, S. & Palva, J. M. Phase transfer entropy: A novel phase-based measure for directed connectivity in networks coupled by oscillatory interactions. NeuroImage 85, 853–872 (2014).

31. Vicente, R., Wibral, M., Lindner, M. & Pipa, G. Transfer entropy—a model-free measure of effective connectivity for the neurosciences. J Comput Neurosci 30, 45–67 (2011).

32. Chang, L. J., Gianaros, P. J., Manuck, S. B., Krishnan, A. & Wager, T. D. A Sensitive and Specific Neural Signature for Picture-Induced Negative Affect. PLoS Biol 13, e1002180 (2015).

33. Zhou, F. et al. A distributed fMRI-based signature for the subjective experience of fear. Nat Commun 12, 6643 (2021).

34. Camacho, M. C. et al. Large-scale encoding of emotion concepts becomes increasingly similar between individuals from childhood to adolescence. Nat Neurosci 26, 1256–1266 (2023).

35. Coutanche, M. N. & Thompson-Schill, S. L. Informational connectivity: identifying synchronized discriminability of multi-voxel patterns across the brain. Front Hum Neurosci 7, 15 (2013).

36. Koster, R. et al. Big-Loop Recurrence within the Hippocampal System Supports Integration of Information across Episodes. Neuron 99, 1342–1354.e6 (2018).

37. Haxby, J. V. et al. Distributed and Overlapping Representations of Faces and Objects in Ventral Temporal Cortex. Science 293, 2425–2430 (2001).

38. Pfeifer, J. H., Iacoboni, M., Mazziotta, J. C. & Dapretto, M. Mirroring others’ emotions relates to empathy and interpersonal competence in children. NeuroImage 39, 2076–2085 (2008).

39. Gallese, V. Mirror neurons and the simulation theory of mind-reading. Trends in Cognitive Sciences 2, 493–501 (1998).

40. Gallese, V., Fadiga, L., Fogassi, L. & Rizzolatti, G. Action recognition in the premotor cortex. Brain 119, 593–609 (1996).

41. Iacoboni, M. Neural mechanisms of imitation. Current Opinion in Neurobiology 15, 632–637 (2005).

42. Nishitani, N. & Hari, R. Temporal dynamics of cortical representation for action. Proc. Natl. Acad. Sci. U.S.A. 97, 913–918 (2000).

43. Rizzolatti, G., Fogassi, L. & Gallese, V. Neurophysiological mechanisms underlying the understanding and imitation of action. Nat Rev Neurosci 2, 661–670 (2001).

44. Meltzoff, A. N. & Marshall, P. J. Human infant imitation as a social survival circuit. Current Opinion in Behavioral Sciences 24, 130–136 (2018).

45. Lee, T.-W., Josephs, O., Dolan, R. J. & Critchley, H. D. Imitating expressions: emotion-specific neural substrates in facial mimicry. Soc Cogn Affect Neurosci 1, 122–135 (2006).

46. Yuan, T., Ji, H., Wang, L. & Jiang, Y. Happy is stronger than sad: Emotional information modulates social attention. Emotion 23, 1061–1074 (2023).

47. Elliot, A. J. & Church, M. A. A hierarchical model of approach and avoidance achievement motivation. Journal of Personality and Social Psychology 72, 218–232 (1997).

48. Shao, X., Li, A., Chen, C., Loftus, E. F. & Zhu, B. Cross-stage neural pattern similarity in the hippocampus predicts false memory derived from post-event inaccurate information. Nat Commun 14, 2299 (2023).

49. Wagner, I. C., Van Buuren, M., Bovy, L. & Fernández, G. Parallel Engagement of Regions Associated with Encoding and Later Retrieval Forms Durable Memories. J. Neurosci. 36, 7985–7995 (2016).

50. De C. Hamilton, A. F. & Grafton, S. T. Action Outcomes Are Represented in Human Inferior Frontoparietal Cortex. Cerebral Cortex 18, 1160–1168 (2008).

51. Carr, L., Iacoboni, M., Dubeau, M.-C., Mazziotta, J. C. & Lenzi, G. L. Neural mechanisms of empathy in humans: A relay from neural systems for imitation to limbic areas. Proc. Natl. Acad. Sci. U.S.A. 100, 5497–5502 (2003).

52. Shamay-Tsoory, S. G., Aharon-Peretz, J. & Perry, D. Two systems for empathy: a double dissociation between emotional and cognitive empathy in inferior frontal gyrus versus ventromedial prefrontal lesions. Brain 132, 617–627 (2009).

53. Fusar-Poli, P. et al. Functional atlas of emotional faces processing: a voxel-based meta-analysis of 105 functional magnetic resonance imaging studies. Journal of psychiatry and neuroscience 34, 418–432 (2009).

54. Kringelbach, M. L. The human orbitofrontal cortex: linking reward to hedonic experience. Nature reviews neuroscience 6, 691–702 (2005).

55. Olszanowski, M., Wróbel, M. & Hess, U. Mimicking and sharing emotions: a re-examination of the link between facial mimicry and emotional contagion. Cognition and Emotion 34, 367–376 (2020).

56. Prochazkova, E. & Kret, M. E. Connecting minds and sharing emotions through mimicry: A neurocognitive model of emotional contagion. Neuroscience & Biobehavioral Reviews 80, 99–114 (2017).

57. Hirsch, J., Zhang, X., Noah, J. A. & Bhattacharya, A. Neural mechanisms for emotional contagion and spontaneous mimicry of live facial expressions. Philosophical Transactions of the Royal Society B: Biological Sciences 378, 20210472 (2023).

58. Ma, X. et al. Own Race Eye-Gaze Bias for All Emotional Faces but Accuracy Bias Only for Sad Expressions. Front. Neurosci. 16, 852484 (2022).

59. Drimalla, H., Baskow, I., Behnia, B., Roepke, S. & Dziobek, I. Imitation and recognition of facial emotions in autism: a computer vision approach. Molecular Autism 12, 27 (2021).

60. Yang, H. et al. Disrupted intrinsic functional brain topology in patients with major depressive disorder. Mol Psychiatry 26, 7363–7371 (2021).

61. Bakdash, J. Z. & Marusich, L. R. Repeated Measures Correlation. Front. Psychol. 8, 456 (2017).

62. Zimeo Morais, G. A., Balardin, J. B. & Sato, J. R. fNIRS Optodes’ Location Decider (fOLD): a toolbox for probe arrangement guided by brain regions-of-interest. Sci Rep 8, 3341 (2018).

63. Chan, M. M. Y. & Han, Y. M. Y. Differential mirror neuron system (MNS) activation during action observation with and without social-emotional components in autism: a meta-analysis of neuroimaging studies. Molecular Autism 11, 72 (2020).

64. Nguyen, T., Miguel, H. O., Condy, E. E., Park, S. & Gandjbakhche, A. Using Functional Connectivity to Examine the Correlation between Mirror Neuron Network and Autistic Traits in a Typically Developing Sample: A fNIRS Study. Brain Sciences 11, 397 (2021).

65. Sun, P.-P. et al. Feasibility of Functional Near-Infrared Spectroscopy (fNIRS) to Investigate the Mirror Neuron System: An Experimental Study in a Real-Life Situation. Front. Hum. Neurosci. 12, 86 (2018).

66. Hou, X. et al. NIRS-KIT: a MATLAB toolbox for both resting-state and task fNIRS data analysis. Neurophotonics 8, 010802 (2021).

67. Fishburn, F. A., Ludlum, R. S., Vaidya, C. J. & Medvedev, A. V. Temporal Derivative Distribution Repair (TDDR): A motion correction method for fNIRS. NeuroImage 184, 171–179 (2019).

68. Pinti, P., Scholkmann, F., Hamilton, A., Burgess, P. & Tachtsidis, I. Current Status and Issues Regarding Pre-processing of fNIRS Neuroimaging Data: An Investigation of Diverse Signal Filtering Methods Within a General Linear Model Framework. Front Hum Neurosci 12, 505 (2019).

69. Hoshi, Y. Functional near-infrared spectroscopy: current status and future prospects. JBO 12, 062106 (2007).

70. Xia, P., Zhang, L. & Li, F. Learning similarity with cosine similarity ensemble. Information Sciences 307, 39–52 (2015).

71. Esteban, O. et al. fMRIPrep: a robust preprocessing pipeline for functional MRI. Nat Methods 16, 111–116 (2019).

72. Gorgolewski, K. et al. Nipype: a flexible, lightweight and extensible neuroimaging data processing framework in python. Front Neuroinform 5, 13 (2011).

73. Gorgolewski, K. J., Nichols, T., Kennedy, D. N., Poline, J.-B. & Poldrack, R. A. Making replication prestigious. Behav Brain Sci 41, e131 (2018).

74. Maldjian, J. A., Laurienti, P. J., Kraft, R. A. & Burdette, J. H. An automated method for neuroanatomic and cytoarchitectonic atlas-based interrogation of fMRI data sets. NeuroImage 19, 1233–1239 (2003).

